# A novel retroviral vector system to analyze expression from mRNA with retained introns using fluorescent proteins and flow cytometry

**DOI:** 10.1101/551846

**Authors:** Patrick E. H. Jackson, Jing Huang, Monika Sharma, Sara K. Rasmussen, Marie-Louise Hammarskjold, David Rekosh

## Abstract

The ability to overcome cellular restrictions that exist for the export and translation of mRNAs with retained introns is a requirement for the replication of retroviruses and also for the expression of many mRNA isoforms transcribed from cellular genes. In some cases, RNA structures have been identified in the mRNA that directly interact with cellular factors to promote the export and expression of isoforms with retained introns. In other cases, a viral protein is also required to act as an adapter. In this report we describe a novel vector system that allows measurement of the ability of *cis*- and *trans*-acting factors to promote the export and translation of mRNA with retained introns.

One reporter vector used in this system is derived from an HIV proviral clone engineered to express two different fluorescent proteins from spliced and unspliced transcripts. The ratio of fluorescent signals is a measurement of the efficiency of export and translation. A second vector utilizes a third fluorescent protein to measure the expression of viral export proteins that interact with some of the export elements. Both vectors can be packaged into viral particles and be used to transduce cells, allowing expression at physiological levels from the integrated vector.

## Introduction

Intron retention is an important form of alternative splicing that is prevalent during retroviral replication, and is also found in the regulation of cellular genes. Special mechanisms have been described that promote the export and translation of mRNAs with retained introns^1^. Simple retroviruses, such as the Mason-Pfizer monkey virus (MPMV), rely on an RNA element present in the transcript that retains an intron, which binds to a cellular protein complex consisting of Nxf1 and the co-factor Nxt1 to promote export and translation^2–4^. This element is referred to as a constitutive transport element (CTE) due to the lack of requirement for viral factors. CTEs have also been found in cellular genes, including within intron 10 of *NXF1* where a CTE functions to export an RNA that is translated into a short form of Nxf1^5–7^.

A more complex retrovirus, HIV-1, requires the nucleo-cytoplasmic export of both unspliced and incompletely spliced viral RNA transcripts for the translation of essential viral proteins and for the packaging of progeny viral genomes^8, 9^. For these mRNAs, export and translation is dependent on the viral Rev protein^10, 11^ and an RNA secondary structure called the Rev Response Element (RRE)^12–14^. Rev binding and multimerization on the RRE permits the assembly of cellular factors, including Crm1 and Ran-GTP, to form an export-competent ribonucleoprotein complex^15, 16^. In contrast, the completely spliced HIV transcripts can be exported and translated in the absence of Rev. Other complex retroviruses, such as equine infectious anemia virus (EIAV) (Rev and RRE)^17^, HTLV (Rex and RexRE)^18^, mouse mammary tumor virus (Rem and RmRE)^19^, and the youngest family of human endogenous retroviruses, HERV-K (Rec and RcRE)^20, 21^, use an analogous mechanism to accomplish the export and translation of intron-containing transcripts.

HIV is notable for the high degree of sequence diversity exhibited during natural infection^22^ and the Rev-RRE system shows significant variation in functional activity between different viral isolates from different hosts^23^, and between isolates from the same host at different time points during infection^24, 25^. While the role of Rev-RRE functional activity differences in HIV pathogenesis has not been fully elucidated, there is evidence that it is an important factor in clinical disease. For example, high RRE activity has been shown to correlate with an increased rate of decline in CD4 count^26, 27^. Conversely, low Rev activity has been associated with prolonged survival in the pre-ART era^28^ and Rev activity has been correlated with the sensitivity of HIV infected T-cells to CTL killing^29^. In experimental infection of ponies with a related virus, EIAV, variation in Rev functional activity was observed during the course of infection, and functional activity differences correlated with clinical disease state^17, 30^.

Variations in the functional activity of the HIV Rev-RRE system have previously been assessed with subgenomic reporter assays^24, 31, 32^ or lentiviral vector packaging assays^23^. Rev-dependent fluorescent reporter systems have also been developed for use in detecting HIV infection, but these have not been used to quantify differences in Rev-RRE activity^33^. Existing functional assays are limited by the multiple steps needed for sample preparation, which often leads to variation between experiments and low throughput. More importantly, nearly all existing assay systems have measured Rev-RRE function using transient transfection of non-lymphoid cell lines.

In order to further correlate the variation that is seen in the Rev-RRE system from different HIV viral isolates with clinical disease states, and to identify additional CTEs present in cellular genes, we developed a new assay system that quickly allows the functional evaluation of large numbers of putative export elements and trans-acting factors. The assay utilizes fluorescent proteins as reporters. A novel aspect of this system is that it uses packageable retroviral constructs, so that after packaging and transduction of target cells, expression can be measured from chromosomally integrated proviral sequences. The data in this report demonstrates the effectiveness of this system in evaluating the expression of mRNA with retained introns mediated by the HIV Rev-RRE axis, by elements from other viruses, and by cellular CTEs.

## Results

### A fluorescence-based high-throughput assay of HIV Rev-RRE functional activity

The HIV provirus is a transcription unit that produces several different mRNA isoforms from a single promoter. Some of these isoforms are unspliced or incompletely spliced, retaining one or more introns, and thus require Rev and the RRE for nucleocytoplasmic export and expression of the proteins they encode. Other isoforms are fully spliced and do not require Rev and the RRE for expression. An assay system to readily measure the functional activity of the HIV Rev-RRE axis was designed by altering the full-length HIV proviral construct NL4-3^34^ to express two fluorescent proteins, one in a Rev-dependent fashion and the other in a Rev-independent fashion (Figure 1). Briefly, the Rev-dependent signal was created by truncating the *gag* gene, deleting the frameshift signal and inserting an in-frame cassette consisting of a *cis*-acting hydrolase element (CHYSEL) sequence^35^ and the *eGFP* gene. The CHYSEL sequence is a self-cleaving 2A peptide derived from a picornavirus that induces ribosome skipping at its C-terminus. This permits the expression of multiple proteins from a single transcript, with the downstream protein (in this case eGFP) being modified by the inclusion of an additional proline at the N-terminus. This alteration would be expected to modify the provirus to express a truncated Gag protein and eGFP from the unspliced Gag-Pol mRNA. As this mRNA contains two introns, it would require Rev and the RRE for export and translation. The Rev-independent signal was created by deleting a portion of *nef* and replacing it with either an *mCherry* or *TagBFP* sequence. Since Nef is made from a fully spliced mRNA that does not require Rev for export and expression, expression of these proteins would not be expected to require Rev. Additional modifications were made to the HIV-derived construct to prevent Rev production from the reporter vector, to further render the construct replication-incompetent, and to permit the easy exchange of RRE sequences within *env*.

**Figure 1.**
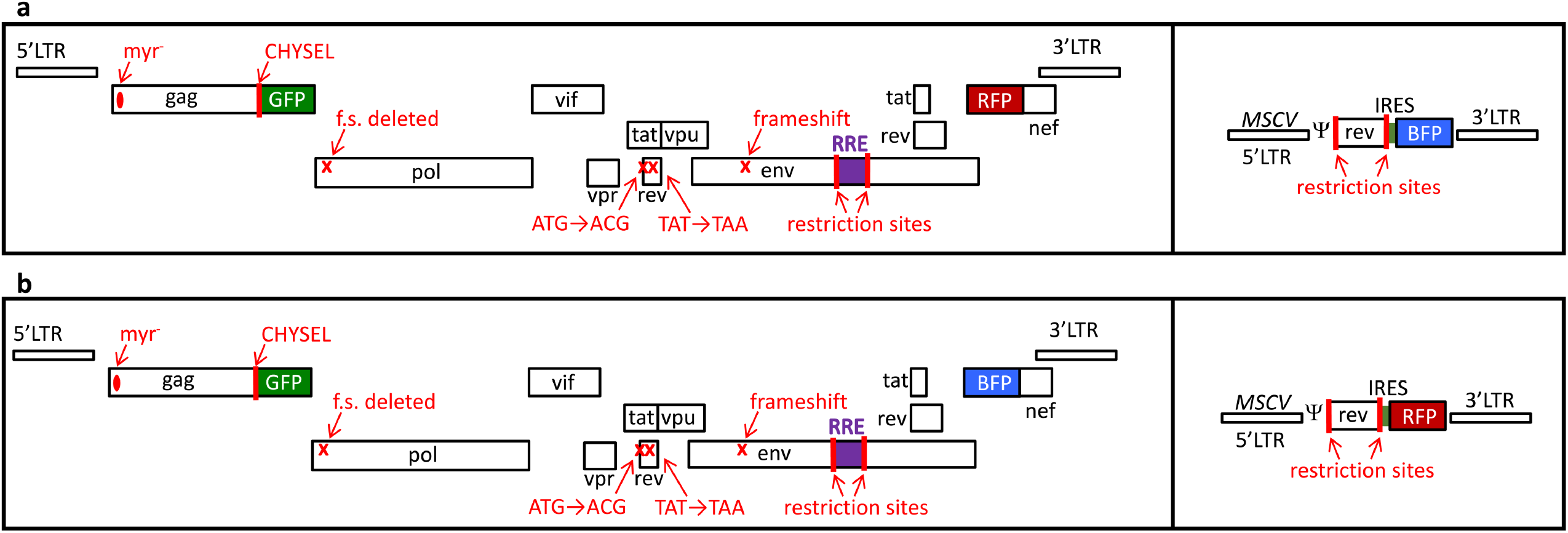
Schematic of retroviral vector constructs used in this study. The two-color constructs (left boxes) are based on the HIV NL4-3 genome with the following modifications from left to right: mutation of the myristoylation signal in *gag*, truncation of *gag*, addition of a CHYSEL sequence and an *eGFP* gene in the *gag* reading frame, deletion of the frameshifting site, missense mutation of the *rev* start codon, insertion of a stop codon in the first exon of rev, introduction of a frameshift mutation in *env*, placement of an XmaI and an XbaI site flanking the RRE within env, and deletion of the beginning of *nef* with the substitution of a second fluorescent protein in this position. The single-color constructs (right boxes) are MSCV vectors modified with a *rev* gene flanked by two restriction enzyme sites as well as an in-frame IRES and a fluorescent protein gene. Two construct sets were used in this study. (a) Construct set A has an eGFP in the *gag* reading frame, mCherry in the *nef* reading frame, and TagBFP in the Rev vector. (b) Construct set B has an eGFP in the *gag* reading frame, TagBFP in the *nef* reading frame, and mCherry in the Rev vector. Myr = myristoylation site, CHYSEL = cis-acting hydrolase element, f.s. = frameshift site, LTR = long terminal repeat, GFP = green fluorescent protein, BFP = blue fluorescent protein, RFP = red fluorescent protein, MSCV = murine stem cell virus, IRES = internal ribosomal entry site, ψ = packaging signal.

Next, we verified that the construct would express TagBFP or mCherry constitutively and eGFP only in the presence of Rev. Both HIV-derived constructs were transfected individually into 293T/17 cells along with varying masses of a plasmid that expressed Rev from the CMV immediate early promoter (pCMV-Rev). Flow cytometry was performed to determine the mean fluorescent intensity of the Rev-dependent signal (eGFP) and the Rev-independent signal (mCherry or TagBFP). Gating was conducted as described in Methods and Figure 2 (a-c). Relative Rev-RRE functional activity was calculated as the ratio of Rev-dependent to Rev-independent signal for each experimental condition.

**Figure 2.**
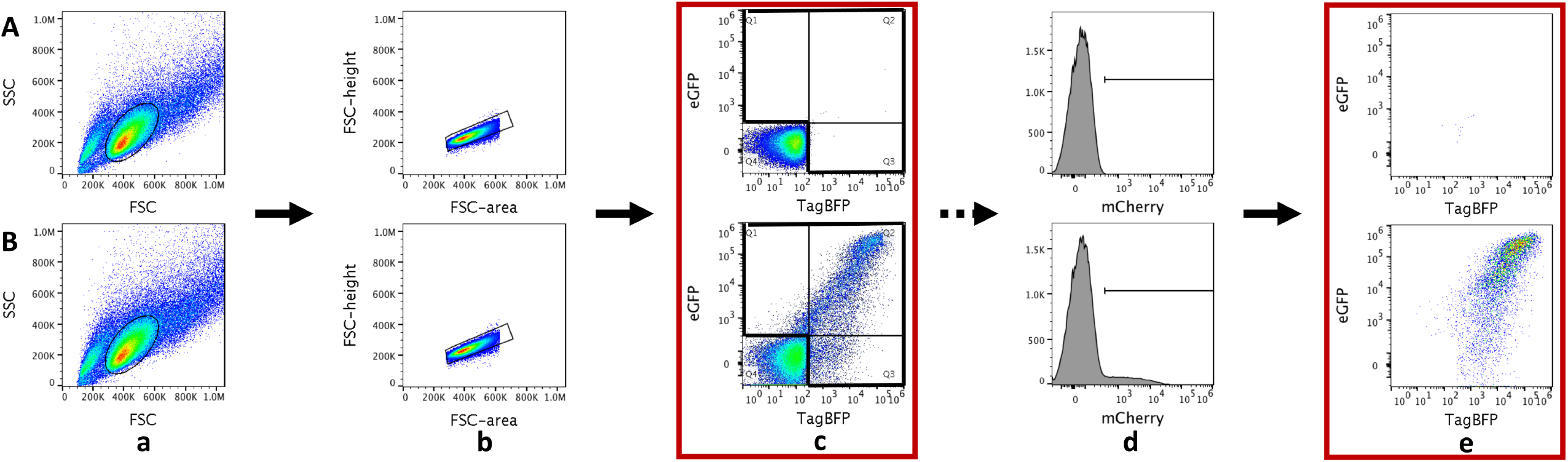
Assay gating strategies. The top panels (A) show untransfected cells. The bottom panels (B) show cells transfected with the vector constructs shown in Figure 1B. 293T/17 cells for analysis were identified by gating first on a forward-scatter versus side-scatter plot (a) followed by an additional gate for single cells constructed on a forward-scatter area versus forward-scatter height plot (b). The fluoresence intensity (eGFP versus TagBFP) of each individual cell is displayed, representing the two markers carried on the HIV-derived construct. (c). Untransfected cells were used to define the double-negative population which was excluded from further analysis. Cells which expressed the HIV-derived construct (i.e. Q1, Q2 and Q3 - eGFP-positive or TagBFP-positive or both) were used for analysis. In other experiments utilizing the MSCV-derived construct carrying a third fluorescent marker, an alternative analysis could be used. In addition to gating on single cells expressing the HIV-derived construct, a gate was created including cells expressing the MSCV-derived construct as well (i.e. mCherry positive) (d). Finally, the population of cells expressing both the HIV-derived construct (i.e. Q1, Q2, or Q3 from c) and the MSCV-derived construct (i.e. mCherry positive from d) was used for analysis (e). Red boxes denote gates used for calculation of functional activity.

Only minimal eGFP signal was observed in the absence of Rev for both HIV-derived constructs (Figure 3). The Rev-RRE activity for both constructs in the 0 ng Rev condition (0.15 for the GFP-mCherry construct and 0.14 for the GFP-BFP construct) was more than 30-fold less than when the lowest tested amount of Rev plasmid (6.25 ng) was included in the transfection (5.59 and 4.33, respectively) and more than 600-fold less than when the maximum tested amount of Rev plasmid (100 ng) was included (value set as 100 for both constructs). While a low level of eGFP “leakiness” is apparent in the flow cytometry dot plots (Figure 3, a and b), this occurred only when there was very high expression of the Rev-independent signal (mCherry or TagBFP) and the system behaved in a highly Rev-responsive fashion. For both constructs, the Rev-independent signal decreased slightly between the 0 ng and 100 ng Rev conditions (2.1 fold for the eGFP-mCherry construct and 2.6 fold for the eGFP-TagBFP construct), but the eGFP signal increased dramatically (313-fold for eGFP-mCherry and 268-fold for eGFP-TagBFP) (Figure S1). The ratio of Rev-dependent to Rev-independent signal was calculated for each condition and both constructs displayed a strongly linear increase in functional activity with increasing amounts of Rev plasmid over the tested range (eGFP-mCherry construct r^2^=0.992, eGFP-TagBFP construct r^2^=0.999) (Figure 3 c).

**Figure 3.**
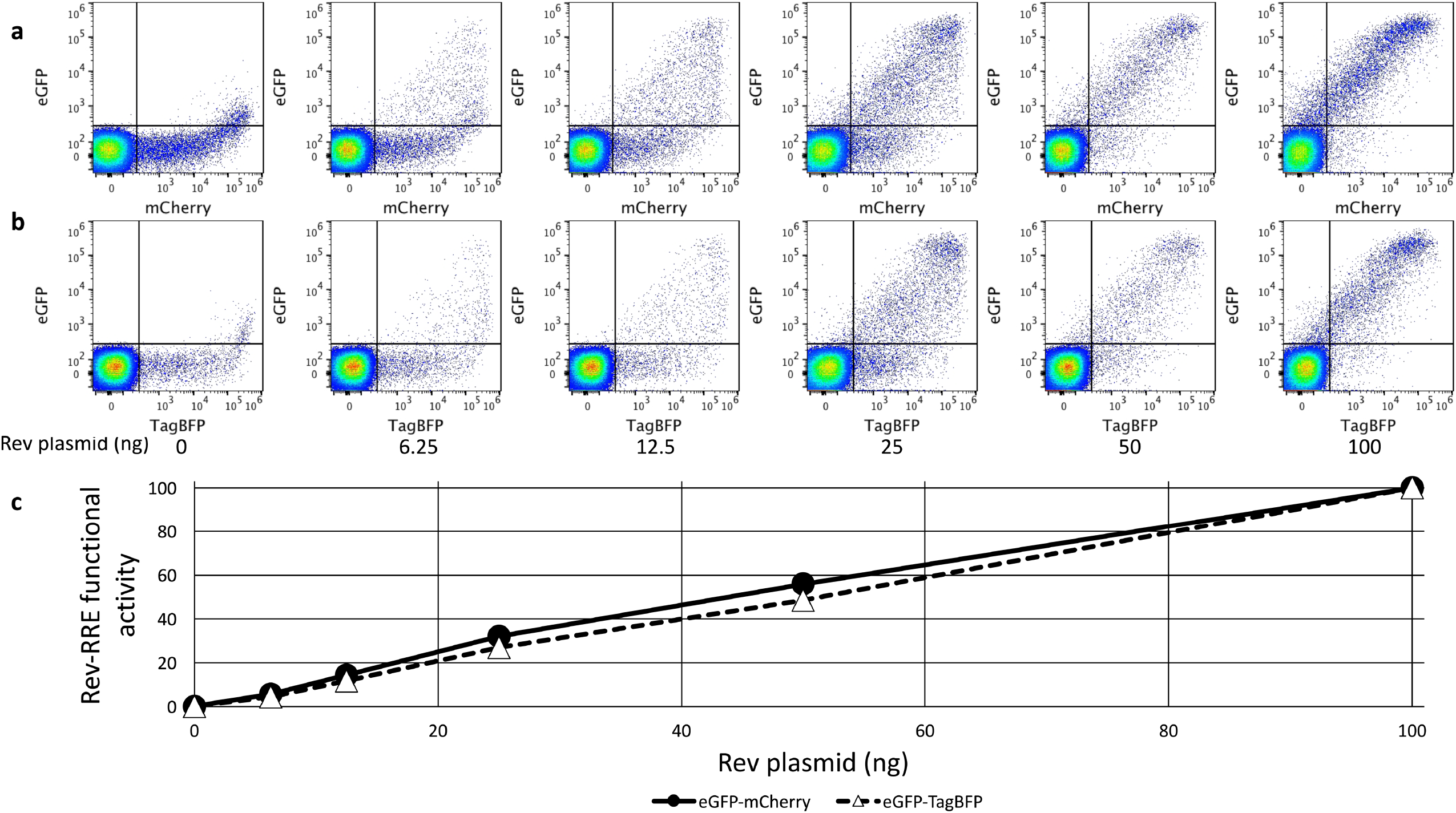
Response of the two color reporter vector to increasing amounts of Rev plasmid. 1000 ng of an eGFP-mCherry (a) or eGFP-TagBFP (b) HIV-derived construct was transfected into 8×10^5^ 293T/17 cells along with a variable amount of CMV-Rev plasmid. Single cells were plotted with Rev-dependent eGFP signal coming from the unspliced mRNA along the y-axis and the Rev-independent signal coming from the fully spliced mRNA along the x-axis (mCherry in a and TagBFP in b). The amount of Rev plasmid transfected along with the two-color construct is listed below each plot. The proportion of the total cells in each plot is shown for every quadrant. Each plot represents at least 100,000 cells (range 107,549 to 119,937 cells). (c) For each construct, the ratio of Rev-dependent (eGFP) to Rev-independent (mCherry or TagBFP) signal MFI was calculated for each experimental condition as a measure of Rev-RRE functional activity (see gating strategy in Figure 2 panel c). The activity in the 100 ng condition for each construct was set as 100. N=2, vertical error bars represent SEM, which are too small to be seen in this figure.

To demonstrate that the CHYSEL sequence functioned to create a free eGFP protein and a separate Gag fragment rather than a Gag-eGFP fusion product, the HIV-derived construct was transfected into 293T/17 cells along with a plasmid that expressed Rev (pCMV-Rev). A Western blot was performed from cell lysate that demonstrated high efficiency functioning of the CHYSEL sequence to produce free eGFP (Figure S2).

### Measurement of differences in Rev and RRE activity

We have previously described Rev and RRE sequences that display significantly different functional activity in prior assays^23, 24^. To assess whether the fluorescence-based assay recapitulates these activity differences, assay constructs were created using previously tested Revs and RREs.

We first tested a known higher-activity RRE (SC3-M57A) along with a known lower-activity RRE (SC3-M0A) sequence that were both derived from naturally occurring viruses in a single patient. These were previously determined to have significantly different activities in a study using a sub-genomic GagPol reporter assay, though both RREs had higher activity than that from NL4-3^24^. New eGFP-TagBFP HIV-derived constructs were created by replacing the native RRE sequence with a 234-nt RRE derived from each virus. The NL4-3 RRE was used for comparison. All three RRE-containing constructs were transfected into 293T/17 cells along with varying amounts of a CMV-Rev plasmid. All three constructs displayed tight Rev responsiveness with minimal calculated activity in the absence of Rev (Figure 4 a). As expected, the SC3-M57A RRE displayed substantially greater functional activity than the SC3-M0A with all tested amounts of Rev plasmid.

**Figure 4.**
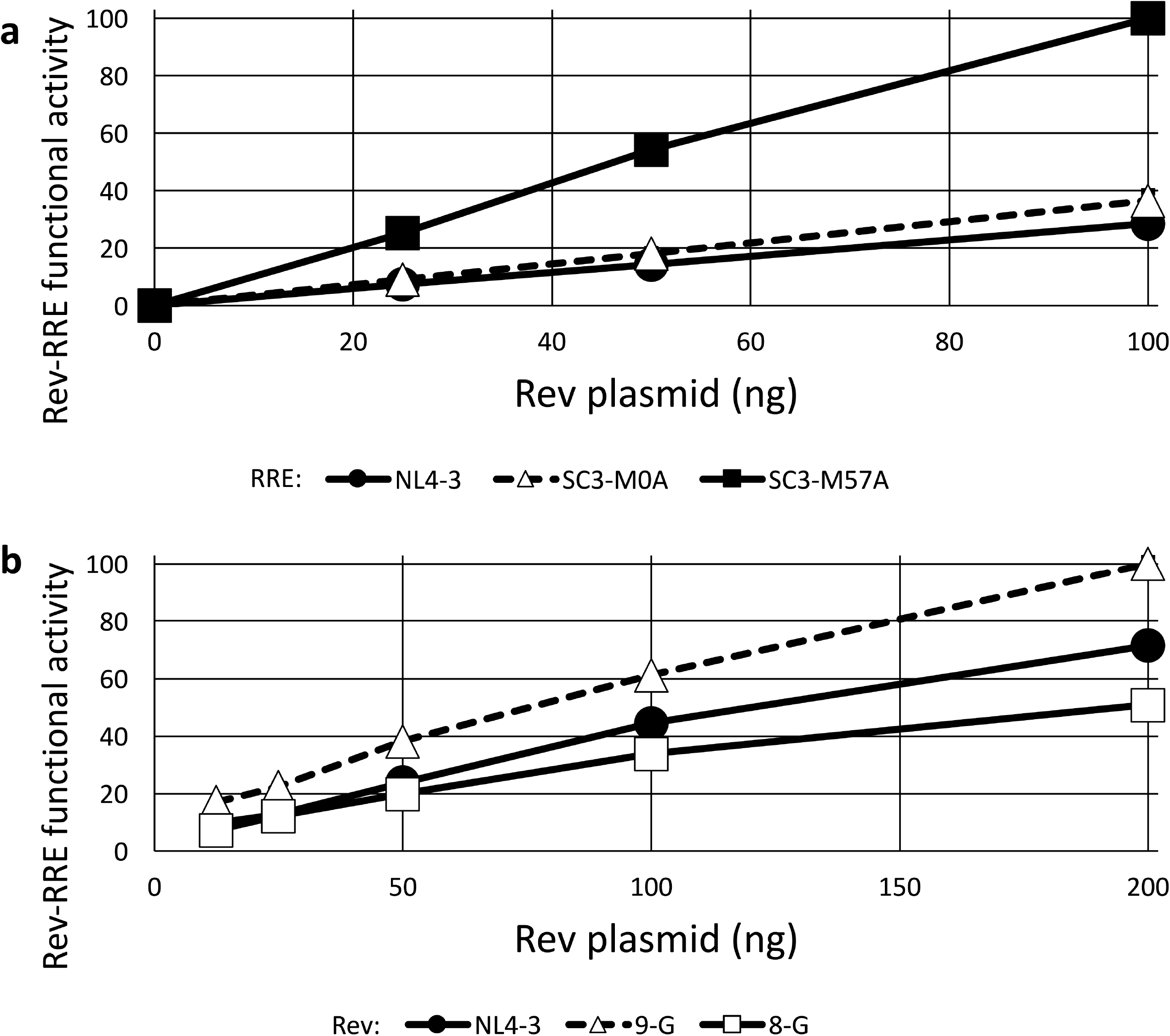
Differential activity of RRE and Rev sequences. (a) Three different 234-nt RREs were cloned into the eGFP/TagBFP HIV-derived construct, including RREs from NL4-3 and two derived from patient isolates (SC3-M0A and SC3-M57A)^24^. 1000 ng of the HIV-derived construct was transfected into 8×10^5^ 293T/17 cells along with 0, 25, 50, or 100 ng of pCMV-SC3-(M0-B/M57-A) Rev^24^. Twenty-four hours after transfection, flow cytometry was performed and the ratio of eGFP/TagBFP signal was determined as a measure of Rev-RRE functional activity. The maximum measured activity was set as 100. (b) Three Rev sequences from NL4-3 or two different primary isolates^54, 55^ were cloned into the MSCV vector carrying an mCherry fluorescent marker. 1000 ng of an eGFP/TagBFP HIV-derived construct containing an NL4-3 RRE sequence was transfected into 8×10^5^ 293T/17 cells along with 12.5, 25, 50, 100, or 200 ng of a pMSCV-Rev-IRES-mCherry construct. Flow cytometry was performed 24 hours after transfection and the data were analyzed as above. N=2, bars, which are too small to be seen in this figure, represent SEM for both graphs.

We also previously described two naturally occurring subtype G HIV-1 Rev proteins with substantially different activity as assessed with a sub-genomic lentiviral vector packaging assay^23^. CMV-Rev plasmids were created expressing high-activity 9-G Rev, low activity 8-G Rev, or NL4-3 Rev. A single eGFP-TagBFP HIV-derived construct containing an NL4-3 RRE was transfected into 293T/17 cells along with varying amounts of each of the CMV-Rev constructs. The fluorescent assay replicated the previously observed difference in activity, with the expected high-activity 9-G Rev yielding substantially greater Rev-RRE functional activity than the 8-G Rev (Figure 4 b). Interestingly, we have previously shown that the difference in 9-G Rev and 8-G Rev functional activity is not correlated with steady state Rev protein levels^23^.

### Evaluation of non-Rev/RRE nuclear export systems

To determine whether the fluorescence-based assay system could detect expression accomplished using mechanisms other than the HIV-1 Rev-RRE system, constructs were created containing elements from the “simple retrovirus” Mason-Pfizer monkey virus (MPMV), the human *NXF1* gene, and the human endogenous retrovirus HERV-K which are known to promote the export and translation of mRNAs with retained introns.

The MPMV CTE directly mediates interaction between the viral mRNA with a retained intron and cellular export factors without the requirement for an additional viral protein. An eGFP-mCherry HIV-derived construct was created with the 162-nt MPMV CTE substituting for the HIV RRE within *env*. A negative control construct was created for comparison by removing the RRE and not substituting an alternative element. Both the MPMV CTE construct and the no element construct were transfected into 293T/17 cells along with plasmids to overexpress the cellular factors NXF1 and NXT1 that have been shown previously to enhance expression of proteins from mRNA whose export has been mediated by the CTE function in 293T cells^36^. The vector with the MPMV CTE showed clear functional activity that was 8-fold higher than background (*P*=3.11×10^−8^) (Figure 5 a).

**Figure 5.**
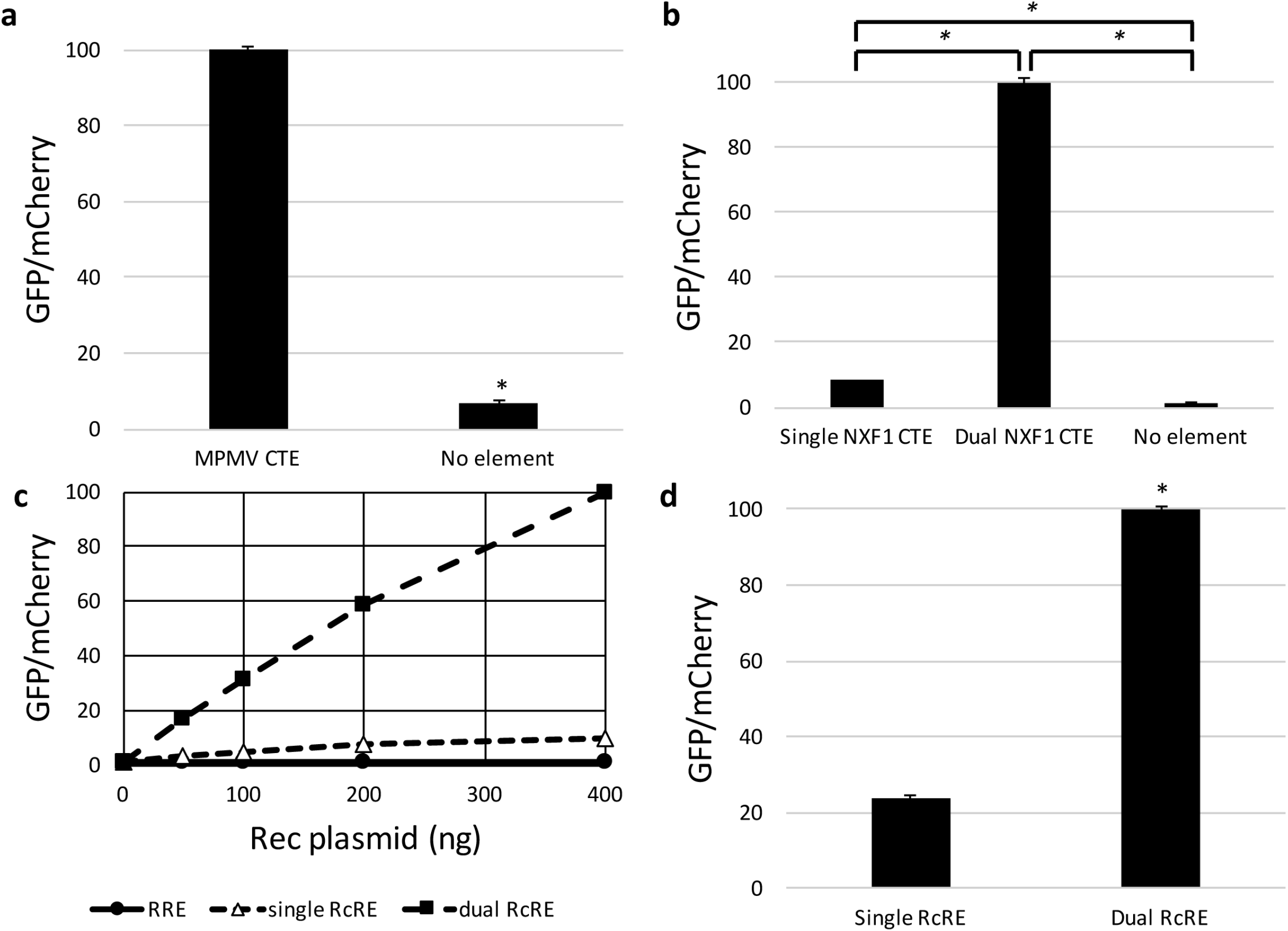
Activity measurements of heterologous viral and cellular RNA export elements in the HIV-derived reporter construct. The eGFP-mCherry HIV-derived construct was modified to carry a variety of RNA transporting elements (or no element) instead of the RRE seen in Figure 1. The modified vectors were then transfected into 293T cells and eGFP and mCherry fluorescence was measured. (a). An HIV-derived construct containing an MPMV CTE was tested along with a construct containing no element. The activity of the MPMV CTE construct was set as 100 and the other values were normalized accordingly. * represents *P*<0.05, N=3, bars represent SEM. (b). HIV-derived constructs containing either one or two copies of the CTE from *NXF1* intron 10 were tested along with a construct containing no element. Activity of the dual *NXF1* CTE construct was set as 100 and the other values were normalized accordingly. In both (a) and (b), Nxf1 and Nxt1 were co-expressed plasmids under the control of the CMV-IE promoter. * represents *P*<0.05, N=2, bars represent SEM. (c) HIV-derived constructs containing single or dual HERV-K RcRE elements or an NL4-3 RRE were co-transfected together with either 0, 50, 100, 200, or 400 ng CMV-Rec plasmid. The activity of the dual RcRE construct in the 400 ng CMV-Rec condition was set as 100 and the other values were normalized accordingly. (d) In a separate experiment, HIV-derived constructs containing single or dual HERV-K RcRE elements were transfected along with 200 ng CMV-Rec plasmid. The activity of the dual RcRE construct was set as 100 and the other values normalized accordingly. * represents *P*<0.05, N=2, bars represent SEM. GFP = green fluorescent protein, MPMV = Mason-Pfizer monkey virus, CTE = constitutive transport element, RcRE= Rec Response Element from HERV-K.

A cellular CTE with a similar sequence and nearly identical structure to one of the 70-nt degenerate repeats present in the MPMV CTE is found in the human NXF1 gene within its intron 10 ^5^. While the 162-nt MPMV CTE contains two internal RNA loops known to interact with cellular export factors, the 96-nt NXF1 CTE contains only a single loop. We created constructs that contained either one or two NXF1 CTEs to assess the functional implication of the number of NXF1 binding loops for vector activity. These constructs were then transfected into 293T/17 cells along with plasmids that overexpress the cellular factors NXF1 and NXT1. The single CTE construct displayed 3-fold increased activity relative to the no element background (*P*=0.012), while the dual CTE construct displayed much greater activity, 33-fold over background (*P*=0.00001) (Figure 5 b). Taken together, these results show that the vector system can be used to assess the function of potential viral and cellular CTEs.

Like other complex retroviruses, the HERV-K (HML-2) provirus contains both an RNA element RRE analogue called RcRE and a Rev viral protein analogue called Rec^37, 38^. To assess whether the activity of this system could be scored in the fluorescent assay, HIV-derived constructs were created in which the RRE was replaced with either one or two 433-nt RcRE sequences in tandem. The RcRE containing HIV-derived constructs, and an RRE-containing construct for reference, were transfected into 293T/17 cells along with varying amounts of a CMV-Rec plasmid. As expected, the RRE-containing construct did not display significant functional activity, as Rec does not function on the RRE^37^ (Figure 5 c). Both the single RcRE and dual RcRE constructs showed linear increases in functional activity with increasing amounts of Rec plasmid (r^2^=0.931 and 0.991, respectively) (Figure 5 c). The dual RcRE construct showed substantially greater activity than the single RcRE construct in the presence of functional Rec (*P*=0.00016) (Figure 5 d). Thus, it should be possible to use this vector to detect functional endogenous Rec expression in cells that have active Herv-K expression.

### Specific inhibition of the Rev-RRE pathway by small molecules

Due to its central role in the HIV life cycle, the Rev-RRE regulatory axis is an attractive therapeutic target. Small molecules that inhibit the Rev-RRE pathway have been developed, though none have yet entered clinical use for the treatment of HIV^39^. A heterocyclic compound designated 103833 and its derivatives have previously been shown to inhibit Rev-RRE functional activity in luciferase and GagPol subgenomic reporter assays^40, 41^. We tested the effect of this compound on both Rev-RRE activity and MPMV CTE activity in the fluorescent assay. 293T/17 cells were pre-treated with varying concentrations of 103833 and the incubation with the compound was continued after transfection with two different sets of plasmids. The effect on the Rev-RRE pathway was tested by transfecting cells with the two color fluorescent construct containing the NL4-3 RRE along with a separate CMV-Rev construct. The effect on the CTE pathway was tested by transfecting cells with the two color fluorescent construct containing the MPMV CTE along with plasmids to overexpress NXF1 and NXT1.

For both constructs, the signal intensity of the fluorescent protein expressed from the *nef* position, mCherry, was not affected by the presence of 103833 (Figure 6 a, boxes). The signal intensity of the protein expressed from the *gag* position, eGFP, showed an inverse relationship with 103833 concentrations only in the case of the RRE-containing construct but not for the CTE-containing construct (Figure 6 a, circles). Thus, the functional activity of the Rev-RRE construct decreased with increasing 103833 concentrations, but the functional activity of the CTE construct remained constant. This is consistent with specific inhibition of the Rev-RRE pathway as previously described. Additionally, the fact that the mCherry signal was unaltered shows that it is a good internal indicator for non-specific effects and toxicity. These results demonstrate the utility of the assay for drug screening.

**Figure 6.**
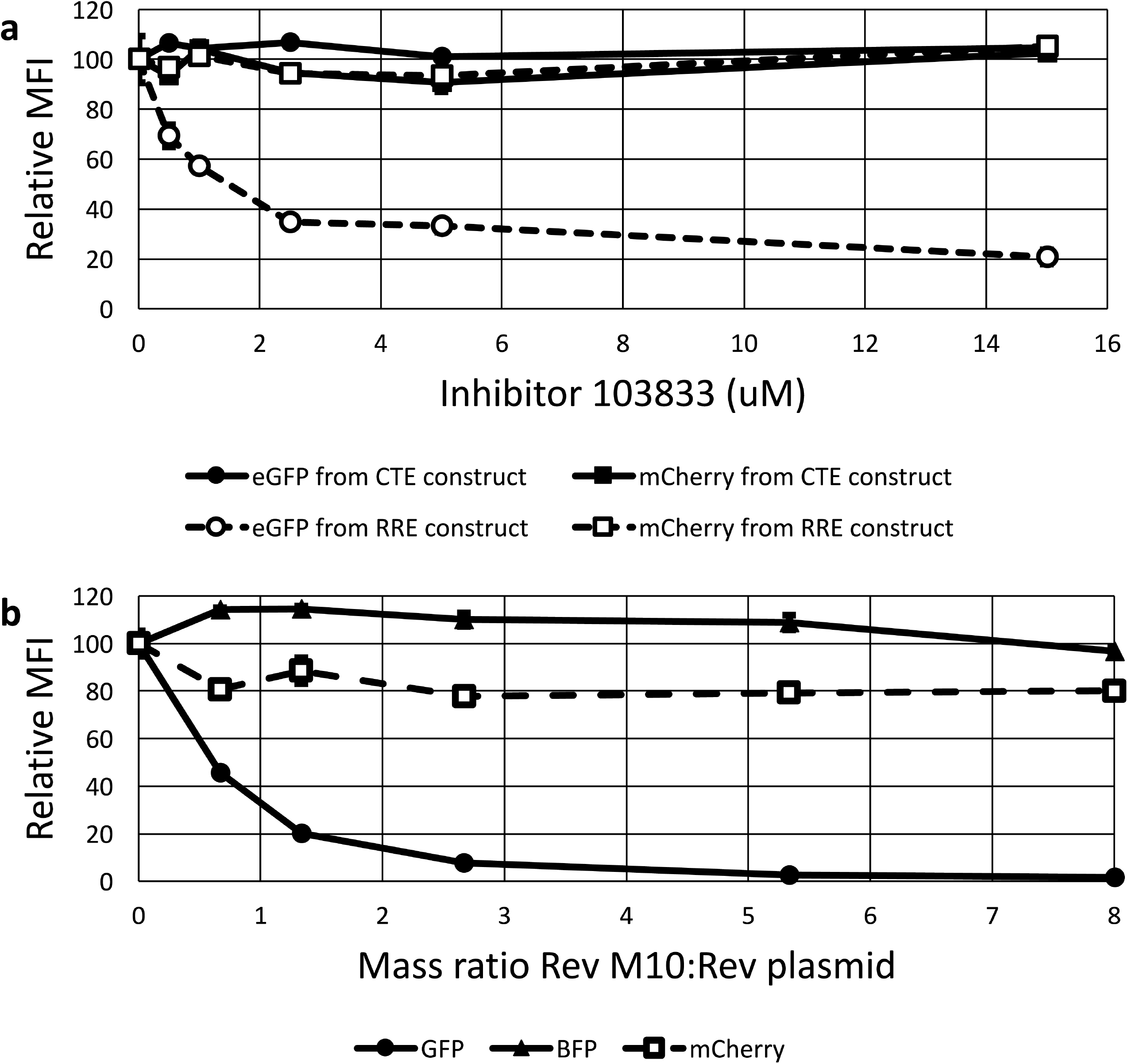
The effect of inhibitors of Rev function on fluorescent protein expression from the reporter vectors. (a) Compound 103833 decreases Rev-RRE functional activity in a dose dependent manner but does not affect CTE function. Two vector constructs were tested in the presence of compound 103833, a Rev-pathway small molecule inhibitor. The RRE construct was identical to that in Figure 1a. The CTE construct was created by replacing the RRE with an MPMV CTE. The constructs were transfected into 293T/17 cells with Rev (RRE construct) or Nxf1 and NxT1 (CTE construct). Cells were pre-treated with the indicated concentrations of 103833 for 24 hours before transfection and remained in the medium until 24 hours after transfection when the cells were harvested for analysis. Both eGFP and mCherry signals were quantified for both constructs. The MFI for each fluorescent signal in the absence of inhibitor was set as 100 and the other values were normalized accordingly. (b) *Trans*-dominant negative Rev M10 inhibits Rev-RRE functional activity in a dose dependent manner. 4×10^5^ 293T/17 cells were transfected with 1000 ng of an eGFP-TagBFP HIV-derived vector along with 75 ng of pMSCV-Rev-IRES-mCherry vector (see Figure 1B). Additionally, a variable mass of a plasmid expressing *trans*-dominant negative Rev-M10 plasmid (pCMV-TD-Rev) was transfected into the system along with an empty vector to equalize total DNA mass of the pCMV plasmids to a total of 600 ng. The MFI of the Rev-dependent (eGFP), Rev-independent (TagBFP), and Rev-linked (mCherry) signals was determined for each ratio of Rev plasmid to Rev-M10 plasmid. For each fluorescent protein, the MFI in the absence of Rev-M10 was set at 100 and the other values were normalized accordingly. N=2, bars, which are too small to be seen in this figure, represent SEM for both graphs. MFI = arithmetic mean fluorescence intensity.

### Rev-RRE pathway inhibition by *trans*-dominant negative Rev M10

An alternative approach to Rev-RRE pathway inhibition is through use of a modified *trans*-dominant negative Rev protein. Rev M10 is a Rev sequence with a mutation in the nuclear export signal and is a well-characterized *trans*-dominant inhibitor of Rev function^42^. We decided to test the effect of Rev M10 on the fluorescence-based functional assay. To do this, 293T/17 cells were transfected with the eGFP-TagBFP HIV-derived construct containing an NL4-3 RRE together with a constant amount of the MSCV construct expressing a bicistronic transcript consisting of the NL4-3 Rev and mCherry genes linked with an internal ribosomal entry site (IRES). The MSCV construct permitted the measurement of mCherry fluorescence as a surrogate for the level of NL4-3 Rev expression. Additionally, the cells were transfected with variable amounts of a plasmid that expressed Rev M10 from a CMV immediate early promoter (pCMV-RevM10). To compensate for the varying amount of Rev M10 plasmid added, an appropriate amount of a pCMV plasmid without an insert was also added.

Figure 6 b demonstrates that as the amount of Rev M10 plasmid added to the transfection increased, the Rev-dependent eGFP signal decreased. Notably, there was no change in the mCherry signal, indicating that the amount of functional NL4-3 Rev remained constant. In addition, the TagBFP signal also remained constant, indicating that Rev-independent expression from the HIV construct was not inhibited. Thus, *trans*-dominant inhibition by Rev M10 could be clearly demonstrated in this assay. This data, combined with the drug inhibition data described above, demonstrates that this assay can easily measure perturbations of Rev-RRE functional activity that are caused by different mechanisms.

### Measurement of Rev-RRE functional activity using packaged assay constructs

Both the HIV-derived RRE-containing constructs and the MSCV-derived Rev-containing constructs were designed to be packageable as viral vectors. The ability to package these constructs allows transduction, rather than transfection of target cells. This permits the assay of Rev and RRE functional activity in different cell types, including primary cells, from an integrated provirus. To demonstrate the ability of the fluorescent assay to assess Rev-RRE functional activity after transduction, a construct carrying the NL4-3 RRE was tested in combination with the three different Revs that were assayed above and known to have varying functional activities.

Three MSCV constructs were created, each carrying a different Rev sequence and expressing a bicistronic transcript encoding both Rev and an mCherry gene separated by an IRES. The Rev sequences included the low-activity (8-G),the high-activity (9-G), and the NL4-3 Revs utilized above^23^. An HIV-derived RRE-containing construct carrying eGFP and TagBFP as markers was used as the fluorescent reporter. The HIV-derived and MSCV-derived constructs were packaged in 293T/17 cells using transient transfection systems with appropriate packaging constructs and pseudotyped using VSV-G. The resulting viral stocks were then used to transduce the CEM-SS lymphoid cell line, either with the HIV-derived construct alone or with the HIV-derived and an MSCV-derived construct in combination.

Cells were successfully transduced with both constructs. Gating was performed on cells expressing the fluorescent marker from the MSCV-derived construct, then an eGFP vs TagBFP plot was constructed to show cells expressing the markers carried on the HIV-derived construct (Figure 7 a). Rev-RRE functional activity (the ratio of eGFP to TagBFP fluorescence) was calculated from the co-transduced population that expressed fluorescent markers from both vectors (i.e. cells included in quadrants 1, 2, and 3 in Figure 7 a). The results showed that the functional activity of NL4-3 Rev and 9-G Rev are both greater than the 8-G Rev (Figure 7d). Thus, the functional activity of the Revs in the T cell line on integrated provirus was similar to that found in the transient transfection assay in 293T/17 cells. Figure 7a-c also shows that a minority of cells that received both the reporter plasmid and the Rev-expressing vector (quadrant Q3) did not express eGFP. The existence of this population would be obscured in assays dependent on bulk protein expression, such as p24 reporter assays, and is only apparent due to the cell-level resolution of this system. Additional studies are underway to evaluate the population of cells that are successfully transduced with both constructs and yet do not express eGFP. Rev function in this population may be compromised in some fashion.

**Figure 7.**
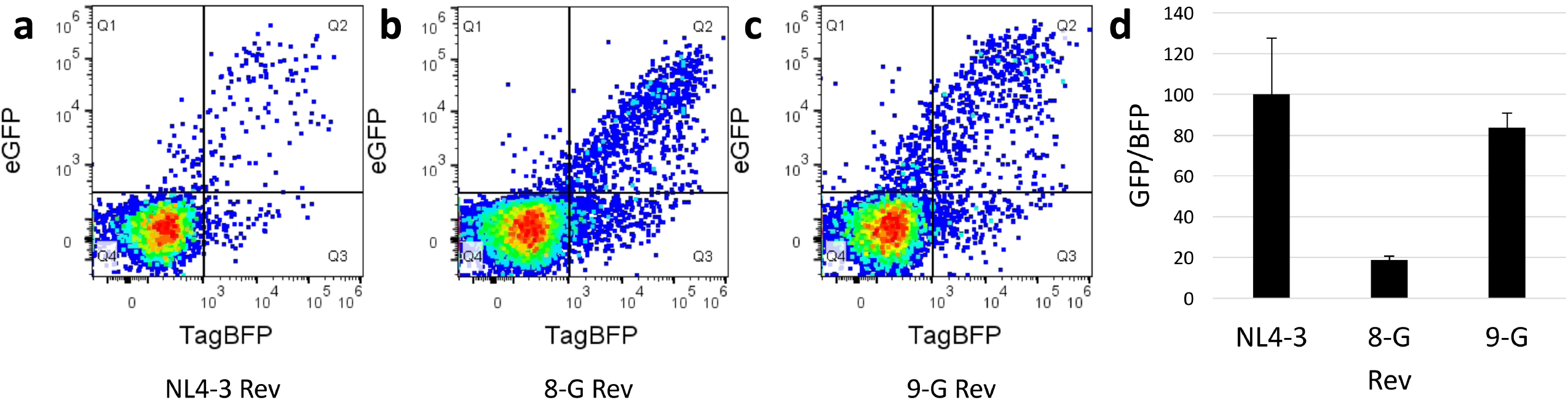
Measurement of Rev-RRE functional activity from an integrated chromatin associated provirus. HIV- and MSCV-derived constructs as in Figure 1 b were packaged and VSV-G pseudotyped in 293T/17 cells. A single HIV-derived construct with an NL4-3 RRE was used along with three MSCV constructs with 9-G, 8-G, or NL4-3 Rev sequences. CEM-SS cells were transduced with both the HIV-derived construct and one of the MSCV-derived constructs at a multiplicity of infection of 0.5. After 72 hours, fluorescence was measured via flow cytometry and gating for analysis was performed as in Figure 2 e (i.e. analyzed cells fluoresce with mCherry, and with eGFP and/or TagBFP). To generate dot plots, single cells were plotted with Rev-dependent eGFP signal along the y-axis and the Rev-independent TagBFP signal along the x-axis with the (a) NL4-3 Rev, (b) 8-G Rev, and (c) 9-G Rev. Flow cytometry plots represent all mCherry-positive single cells measured in three experimental replicates prior to analysis gating. Plot (a) shows 5,690 cells, (b) shows 12,734 cells, and (c) shows 8,534 cells. (D) Relative Rev-RRE functional activity was calculated for each Rev construct as the ratio of eGFP:TagBFP MFI in the set of cells expressing fluorescent markers from both transducing constructs. The 9-G Rev activity was set as 100 and the other values were normalized accordingly. N=3, bars represent SEM.

### Creation of a stable T-cell line containing a single integrated copy of the HIV-derived reporter vector

To establish clonal stable cell lines, CEM-SS cells were transduced with the HIV-derived vector containing mCherry in the *nef* position and eGFP in *gag*. Individual clones expressing mCherry were then derived from the transduced cells by limiting dilution. Clones were analyzed for mCherry and eGFP expression. Clone 5 showed a very uniform and strong mCherry expression with virtually no eGFP or tag-BFP signal (Figure 8, left side plots). Integration of a single copy of the HIV-derived construct within an intergenic region of chromosome 9 (downstream of gene RGP1 and upstream of long non-coding RNA AL1333410; chromosome 9: 35765800) was determined by inverse PCR (data not shown). The cells were then transduced with the MSCV vector expressing NL4-3 Rev and TagBFP (Figure 8, right side plots). About 8% of cells expressed TagBFP (Figure 8 a), demonstrating successful transduction with the MSCV construct, and a similar percent expressed eGFP from the unspliced *gag* transcript (Figure 8 b). It is likely that the different levels of eGFP and tag-BFP expression stem from the fact that the integration site of the MSCV vector in each transduced cell is different, and this leads to different levels of expression from the integrated proviral vector. Significantly there was a good linear correlation of BFP versus eGFP expression levels, indicating that cells that expressed higher Rev levels also expressed higher levels of the mRNA with the retained intron (Figure 8 c). These cells were subjected to cell sorting, allowing us to obtain cells that have different levels of Rev expression that can be used for further studies.

**Figure 8.**
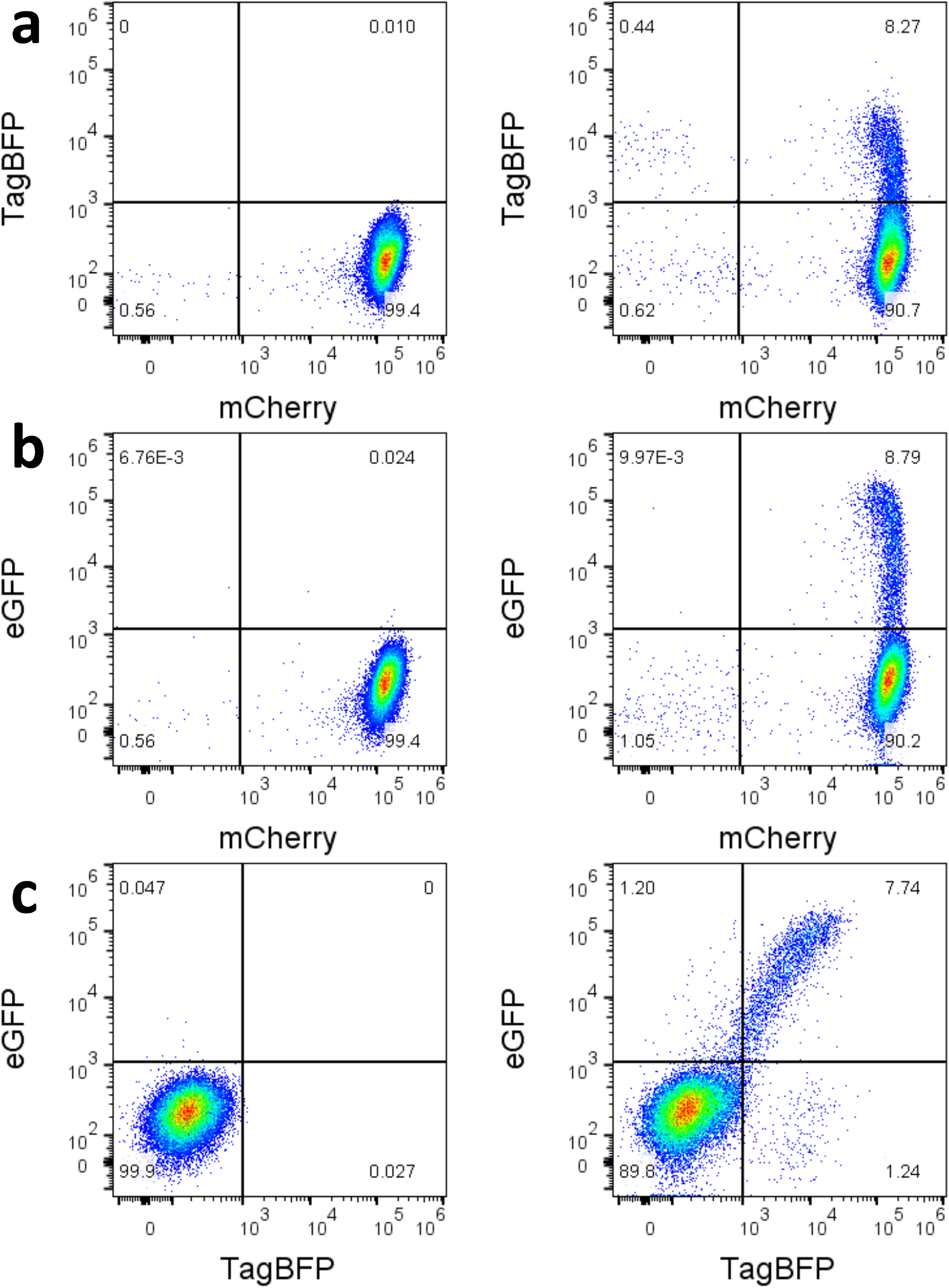
Induction of eGFP expression from a CEM-SS cell line containing a stably integrated single copy of the reporter plasmid. CEM-SS cells were transduced with an eGFP-mCherry HIV-derived construct and a clonal cell line was created by limiting dilution. Thus, almost all cells show a uniformity of mCherry expression (Plot a-left). The cell line was subsequently transduced by a pMSCV-Rev-IRES-TagBFP and fluorescence was measured for all three markers. Untransduced cells are shown on the left, cells transduced with the MSCV-derived Rev/BFP expressing vector are shown on the right. Plot a (right) shows that transduction caused about 8.27% of the cells to express TagBFP. Plot b (right) shows that transduction caused about 8.79% of the cells to express eGFP. Plot c (right) shows that there is a linear relationship between the BFP and eGFP signal, indicating a strong correlation between Rev levels in a cell (BFP) and expression of the unspliced mRNA isoform (eGFP).

## Discussion

In this paper, we describe a novel easily manipulated system that allows the detection and characterization of RNA elements and trans-acting protein factors that promote the nucleocytoplasmic export and translation of mRNAs with retained introns. Intron retention is essential for retrovirus replication. Furthermore, intron retention in cellular genes is increasingly recognized as playing an important role in development and differentiation, as well as in disease states such as cancer and responses to cellular insults^1^. In the case of complex retroviruses, such as HIV and EIAV, there is also considerable evidence that different viral isolates vary significantly in their ability to express intron-containing viral RNA and that these differences are important in pathogenesis and possibly also in the regulation of viral latency^17, 23, 24, 27, 30^. The ability to detect and functionally characterize the RNA elements and proteins involved in the export and translation of mRNAs with retained introns is essential for further insights into the fate and regulation of these mRNAs.

Our assay system improves on previous reporter assays in several important respects. First, the use of three fluorescent proteins as reporters allows the simultaneous measurement of expression from the mRNA with the retained intron, the overall transcription level of the reporter vector, and the level of expression of the trans-acting protein involved in RNA export. Furthermore, the use of flow cytometry to quantify expression levels provides for more rapid acquisition of data compared with prior assays. Many previous reporter assays have required collection of cells and subsequent quantification of non-fluorescent proteins such as p24 by ELISA^24^ or enzymatic activity^32^. In other cases, RNA export function has been measured by the determination of vector titer^23^. The assay described here will permit more rapid determination of the activity of larger numbers of Rev and RRE sequences derived from HIV patients. This will allow the evaluation of the larger data sets that are needed to make good statistical correlations of Rev-RRE activity with specific clinical outcomes. It will also more readily allow the identification of novel RNA elements in cellular genes, and facilitate drug discovery targeting the interaction of RNA signals with the protein factors that recognize them.

Second, nearly all previous reporter assays have measured the expression of mRNA with retained introns from DNA plasmids that have been transiently expressed, despite the fact that there is considerable evidence that mRNA export factors first encounter the mRNA when they are in the process of being newly transcribed from chromatin^43, 44^. In the assay described here, the vectors can be packaged into infectious retrovirus-vector particles which integrate into the host cell genome. This allows measurement of expression of mRNAs with retained introns that have been transcribed from chromatin, rather than transiently from transfected plasmids, and is thus less artificial than previous assays. Overexpression of both the reporter mRNA and export factor can also be avoided by manipulating the multiplicity of infection such that single copies of reporter construct and the gene for the trans-acting protein factor are present in a cell. This is important, as we know, for example, that in the case of HIV high levels of Rev protein expression produce artifactual results and only low levels of Rev are expressed in an infected cell^45^. The ability to package and pseudotype the vector constructs also permits the use of the assay in diverse cell types and primary cells, rather than in only easily transfectable cell lines. Thus, our assay system much more closely resembles the conditions of natural HIV infection and the expression of normal levels of cellular mRNA.

Third, the ability to stably integrate assay constructs into the host cell chromosome also permits the creation of cell lines containing one or both assay constructs. Using the CEM-SS T-cell line, we created a clonal population containing a single integrated copy of an HIV-derived assay construct within an intergenic region. This cell line constitutively expresses the Rev-independent fluorescent marker (mCherry), but expresses the Rev-dependent marker (eGFP) only after transfection or transduction with a Rev-containing construct. When this cell line is transduced with an MSCV-derived construct carrying HIV Rev and TagBFP, eGFP expression increases linearly with TagBFP expression. This demonstrates a close correlation between Rev levels and eGFP expression, and the linear responsiveness of the system. Thus, the creation of this cell line and others will permit more rapid drug screening studies and studies of the cellular determinants involved in the export and translation of mRNA with retained introns during HIV infection. Similar cell lines can also be created using these constructs containing cellular CTEs to evaluate the role of cellular proteins in the export and translation of cellular mRNAs with retained introns.

In summary, we have described a novel high-throughput fluorescence-based assay of intron retention that permits the functional characterization of mechanisms that promote expression from mRNAs with retained introns, including systems from complex and simple retroviruses and cellular genes. This assay represents a substantial improvement from previous assays in that studies can be performed at the level of integrated proviral constructs in a variety of cell lines and primary cell types. Applications for this assay include drug screening for compounds that inhibit or promote the export of mRNAs with retained introns, identification of cellular CTEs, and investigation of the evolution of the Rev-RRE axis in clinical disease progression during HIV infection.

## Methods

### Creation of assay constructs

HIV-derived constructs were created by modifying a plasmid containing the full-length genome of laboratory strain NL4-3 (Genbank accession U26942) (Figure 1)^34, 46^. To create the fluorescent signal from the unspliced transcript, *gag* was truncated at amino acid 407 and a cassette was inserted in-frame containing a CHYSEL sequence derived from porcine teschovirus-1^35^ and the *eGFP* gene^47^. To create the fluorescent signal from the completely spliced transcript, *nef* was deleted and replaced with either an *mCherry*^48^ or *TagBFP* gene^49^. The construct was further modified to silence the native *rev* and to further render the construct replication incompetent^50^. Finally, the 351-nt RRE sequence within *env* was flanked by an XmaI and XbaI site to permit easy replacement.

To create the MSCV constructs, Rev sequences were cloned into MSCV-based plasmids containing the genes for fluorescent proteins (Figure 1). *Rev* was cloned upstream of an IRES-fluorescent protein cassette, such that both Rev and the fluorescent protein are expressed from a bicistronic transcript. Additional non-MSCV Rev-containing plasmids were created using an immediate early CMV promoter to drive Rev expression without a fluorescent marker. Additional descriptions of the assay constructs are available in Supplemental Methods.

Versions of the HIV-derived construct containing different export elements were created by replacing the RRE with alternative sequences. These constructs are denominated in the text as pNL4-3(unspliced transcript fluorescent marker)(transport element)(spliced transcript fluorescent marker), as in pNL4-3(eGFP)(NL4-3 RRE)(mCherry). Similarly, different Rev sequences or other *trans*-acting proteins were substituted in the MSCV-derived construct. These constructs are denominated pMSCV - *trans*-acting protein – IRES – fluorescent marker, as in pMSCV-NL4-3 Rev-IRES-TagBFP. The CMV constructs are denominated as in CMV-NL4-3 Rev.

### Western blot of Gag-CHYSEL-eGFP product

A Western blot was performed to test the cleavage efficiency of the CHYSEL sequence incorporated in the *gag*-CHYSEL-*eGFP* cassette. 293T/17 cells were transfected with constructs expressing eGFP alone, the *gag*-CHYSEL-*eGFP* cassette, a *gag-eGFP* fusion sequence, and mCherry as a negative control. Western blotting was performed on cell lysate using primary antibodies to eGFP and p24^51^. Additional details are available in the Supplemental Methods.

### Transfection-based functional assays

The assay system permits determination of Rev-RRE functional activity at the level of transfection. 293T/17 cells were co-transfected with an HIV-derived construct and either a MSCV-derived construct or CMV-Rev using the polyethylenamine method^52^. Cells were incubated for 24 hours after transfection. Flow cytometry was performed using the Attune NxT flow cytometer with autosampler attachment (Thermo Fischer Scientific). Data analysis, including color compensation, was performed using FlowJo v10 (FlowJo, LLC). Further details are available in the Supplemental Methods.

The gating schemes used are shown in Figure 2. Cells were selected (panel a) and cell aggregates excluded (b). Cells expressing the HIV-derived construct (bottom) were identified by reference to a negative control (top) (c). Untransfected cells in the bottom left quadrant were excluded from further analysis. When an MSCV-derived construct was used, a derived gate from (b) was constructed to include only cells expressing the third fluorescent marker linked to the *trans*-acting protein (d). Finally, cells expressing both constructs were included in a final analysis (e).

To calculate the activity of the intron export mechanism, cells in the appropriate gate for analysis (panel c excluding the bottom left quadrant or e) were considered. The arithmetic mean fluorescence intensity (MFI) was determined for both fluorescent signals of the HIV-derived construct, and the ratio of the unspliced transcript marker MFI (i.e. eGFP) to the spliced transcript marker MFI (e.g. TagBFP or mCherry) was calculated. The resulting ratio was adjusted by subtracting the activity of the 0 ng Rev condition (or equivalent) to compensate for eGFP “leakiness.” Finally, functional activity values within the experiment were expressed as a proportion of the maximum for any condition, which is set as 100.

### Rev titration assay

The responsiveness of the HIV-derived construct to increasing amounts of Rev plasmid was determined. 1000 ng of the HIV-derived construct pNL4-3(eGFP)(NL4-3 RRE)(TagBFP) (pHR5564) was transfected into 8×10^5^ 293T/17 cells along with 0, 6.25, 12.5, 25, 50, or 100 ng of CMV-NL4-3 Rev plasmid (pHR5186). The total mass of DNA in each transfection was kept constant by the addition of varying masses of an empty CMV plasmid (pHR16). Flow cytometry was performed 36 hours after transfection.

### Assay of different RRE sequences

Three HIV-derived constructs expressing eGFP and TagBFP were created by replacing the native RRE sequence with alternative 234-nt RREs. The RREs were from NL4-3 (pHR5580) as well as two patient-derived viruses: SC3-M0A and SC3-M57A (Genbank accession KF559160.1 and KF559162.1; pHR5584 and pHR5586)^24^. In a previous assay system, SC3-M57A was found to have higher activity than SC3-M0A^24, 53^. The constructs were tested individually by transfecting them into 293T/17 cells along with 0, 25, 50, or 100 ng of CMV-SC3 Rev plasmid using a Rev sequence from the same patient as the RRE sequences (Genbank accession KF559146; pHR4776)^24^. The total DNA mass in each condition was kept constant by the addition of a variable mass of an empty CMV construct (pHR16).

### Assay of different Rev sequences

Three MSCV-derived constructs expressing mCherry and one of three different Revs were created. The Rev sequences used were from NL4-3 (pHR5625) and two subtype G HIV viruses: 8-G and 9-G (Genbank accession FJ389367^54^ and JX140676^55^; pHR5629 and pHR5627). In a previous assay system, 9-G was found to have higher activity than 8-G Rev^23^. 293T/17 cells were transfected with 1000 ng pNL4-3(eGFP)(NL4-3 RRE)(TagBFP) (pHR5580) along with 0, 12.5, 25, 50, 100, or 200 ng of each of the MSCV constructs. The total DNA mass in each transfection was kept constant by addition of a variable mass of an empty CMV plasmid (pHR16).

### Assay of HERV-K RcRE and Rec activity

An HIV-derived eGFP-mCherry construct was modified by replacing the HIV RRE sequence with a 433-nt HERV-K RcRE sequence (Genbank accession AF179225.1^37^, pHR5476). An additional HIV construct was created by inserting a second identical RcRE sequence in tandem (pHR5631). Both RcRE-containing constructs and an HIV-derived RRE-containing construct (pHR5460), were tested by transfecting them individually into 293T/17 cells along with 0, 50, 100, 200, and 400 ng of a CMV plasmid containing the HERV-K Rec gene (Genbank accession AY395522.1^38^), CMV-Rec (pHR5313). Flow cytometry was performed 48 hours after transfection. This experiment was repeated for the 200 ng CMV-Rec condition to obtain additional replicates.

Statistical analysis was performed using SPSS v25 (IBM). R-squared values for Figure 5c were calculated using the linear regression function and *P* values for Figure 5d were calculated using an independent-samples T-test.

### Assay of CTE activity

A 96-nt CTE derived from intron 10 of *NXF1* was incorporated into an eGFP-mCherry HIV-derived construct in place of the RRE (see Genbank reference sequence NM_001081491.1 bases 1928 to 2023, pHR5781)^5^. An additional construct was created from this vector by inserting a second *NXF1* CTE within *env* at the NheI site (pHR5783). Another HIV-derived construct was created by replacing the RRE with a single copy of a 162-nt CTE from MPMV (Genbank accession AF033815.1 bases 7388 to 7549, pHR5608)^2, 3^.

Each of the resulting CTE constructs were tested by transfecting 1000 ng of each individually into 293T/17 cells along with CMV-Nxf1 plasmid (pHR3704) and CMV-Nxt1 plasmid (pHR2415) to overexpress these cellular factors. The MPMV CTE experiments were performed using 100 ng Nxf1 plasmid and 200 ng Nxt1; the NXF1 CTE experiments were performed using 100 and 25 ng, respectively. Flow cytometry was performed 48 hours after transfection.

Statistical analysis of the activity differences of the CTE constructs was performed using SPSS Statistics v25 (IBM). P-values were calculated using an independent-samples T-test for Figure 5a and a two-tailed one-way ANOVA with Tukey’s HSD test for Figure 5b.

### Small molecule Rev-RRE pathway inhibitor assay

A previously described heterocyclic compound, 3-amino-5-ethyl-4,6-dimethylthieno[2,3-b]pyridine-2-carboxamide (103833), inhibits Rev-RRE function^40^. 293T/17 cells were pre-treated with varying concentrations of 103833 and the incubation with the compound was continued after transfection. To test the response of the Rev-RRE system to the inhibitor, cells were transfected with 1000 ng of pNL4-3(eGFP)(NL4-3 RRE)(mCherry) (pHR5460) and 100 ng of pMSCV-NL4-3 Rev-IRES-TagBFP (pHR5420). To test the response of the CTE system, cells were transfected with 1000 ng of pNL4-3(eGFP)(MPMV CTE)(mCherry) (pHR5608), 133 ng CMV-Nxf1 (pHR3704), and 33 ng CMV-NxT1 (pHR2415). Data analysis was conducted by gating on cells successfully transduced with the HIV-derived two-color constructs (see Figure 2 panel c). Additional details are available in the Supplemental Methods.

### *Trans*-dominant negative Rev M10 assay

To measure the effect of the co-expression of Rev M10^42^ on Rev-RRE activity, 4×10^5^ 293T/17 cells were transfected with 1000 ng pNL4-3(eGFP)(NL4-3 RRE)(TagBFP) (pHR5580), 75 ng pMSCV-NL4-3 Rev-IRES-mCherry (pHR5625), and either 600 ng, 400 ng, 200 ng, 100 ng, 50 ng, or 0 ng of CMV-Rev M10 (pHR636). Variable amounts of an empty CMV plasmid (pHR16) were included to maintain a constant DNA mass in each transfection. Data analysis was performed by gating on cells successfully transduced with the HIV-derived construct (see Figure 2 panel c). The arithmetic MFI of mCherry was determined to measure relative expression of NL4-3 Rev.

### Packaging and titering HIV and MSCV constructs

The HIV-derived construct pNL4-3(eGFP)(NL4-3 RRE)(TagBFP) (pHR5580) was packaged and VSV-G pseudotyped using a transient second generation packaging system including psPAX2 (Addgene plasmid 12260, pHR5691) and pMD2.G (Addgene plasmid 12259, pHR5693). Three MSCV constructs of the form pMSCV-Rev-IRES-mCherry carrying the NL4-3, 8-G, or 9-G Revs (pHR5625, 5629, 5627 respectively) were packaged as well in a transfection system that included pHIT60-CMV-GagPol^56^ (pHR1854) and pMD2.G (pHR5693). Vector stock titer was determined in 293T/17 cells using constitutively expressed fluorescent proteins as the marker of transduction. Additional details are available in the Supplemental Methods.

### Assay of Rev activity in transduced cells

The relative functional activity of Rev-RRE pairs was assayed by co-transducing CEM-SS cells with viral vectors derived from pNL4-3(eGFP)(NLR-3 RRE)(TagBFP) and the different pMSCV-Rev-IRES-mCherry constructs^57, 58^. As controls, the vectors were also added to the CEM-SS cultures individually. Seventy-two hours after transduction, cells were harvested and flow cytometry was performed. Functional activity was determined by gating on cells transduced with both constructs (see Figure 2 panel e). See the Supplemental Methods for additional details.

### Creation of T-cell line containing the integrated two color reporter construct

To create a cell line with a stably integrated HIV-derived proviral construct carrying eGFP and mCherry, CEM-SS cells were transduced with a VSV-G pseudotyped pNL4-3(eGFP)(NL4-3 RRE)(mCherry) viral vector (pHR5604). A limiting dilution was performed in 96 well plates to generate monoclonal populations, and clones expressing the HIV-derived construct were identified by looking for mCherry expression via fluorescent microscopy. The presence of a single integrated copy of the HIV construct was confirmed by a modified inverse PCR procedure^59^. Additional details are available in the Supplemental Methods.

After identifying a clone with a single integrated HIV-derived construct, intron export was demonstrated by transducing cells with the VSV-G pseudotyped pMSCV-NL4-3 Rev-IRES-TagBFP (pHR5420) construct as above. Seventy-two hours after transduction, eGFP, mCherry, and TagBFP expression was quantified by flow cytometry.

## Data Availability

No datasets were generated or analyzed during the current study. Sequences of the evaluated intron-retention elements are available from Genbank at the accession numbers included in the text. Plasmids and cell lines utilized in this study are available from the authors upon request.

## Acknowledgements

The pMSCV-IRES-mCherry and pMSCV-IRES-TagBFP vectors were gifts from Dario Vignali obtained through Addgene (unpublished). The plasmids psPAX2 and pMD2.G were gifts from Didier Trono obtained through Addgene. Technical assistance and guidance on data interpretation was provided by the University of Virginia Flow Cytometry Core Facility. The p24 hybridoma (183-H12-5C) (Cat# 1513) was obtained through the NIH AIDS Reagent Program, Division of AIDS, NIAID, NIH from Dr. Bruce Chesebro. This work was supported by grants GM 110009, CA206275, AI134208, AI087505 and AI068501 from the National Institutes of Health (NIH) to M-L.H and D.R. P. E. H. J. was supported by grant K08AI136671 from the National Institutes of Health. Salary support for M.-L.H. and D.R. was provided by the Charles H. Ross, Jr., and Myles H. Thaler Endowments at the University of Virginia.

## Author Contributions

P. E. H. J wrote the first draft of the manuscript text, prepared all figures, and performed many of the experiments. M. S. and S. R. performed the experiments relating to HERV-K Rec and the RcRE. J. H. was involved in the development of the two color vector and performed many of the initial experiments not reported here to characterize the system, as well as some of the experiments reported here. D. R. and M.-L. H. conceptualized the assay system, contributed to study design, manuscript revisions, and project oversight. All authors reviewed the manuscript.

