## Supplemental Methods and Figs. S1 and S2 for "A novel retroviral vector system to analyze expression from mRNA with retained introns using fluorescent proteins and flow cytometry"

*Creation of assay constructs.*

Additional modifications were made to the HIV-derived construct to render it unable to replicate and to silence native Rev expression. These modifications are: (a) the second codon of *gag* was mutated from GGT (glycine) to GCT (alanine) to eliminate the myristoylation site, (b) the *rev* start codon was changed to ACG without modifying the amino acid sequence of Tat, (c) an in-frame stop codon was introduced into the first exon of *rev* at amino acid position 23, and (d) a frameshift was introduced into *env* upstream of the RRE.

The MSCV single-color constructs were derived from pMSCV-IRES-mCherry FP (Addgene plasmid # 52114) (pHR5301) and pMSCV-IRES-Blue FP (Addgene plasmid # 52115) (pHR5302), both of which have a multiple cloning site upstream of the IRES and the fluorescent marker.

*Western blot of Gag-CHYSEL-eGFP product.*

A 10 cm plate containing a total of 5 x $10^5$ 293T/17 cells was transfected with (a) 15 ug of a lentiviral vector with a PGK-eGFP cassette (pHR5430), (b) an HIV construct with *gag*-CHYSEL-*eGFP* similar to Figure 1a (pHR5460), (c) an HIV construct with *gag-eGFP* and no intervening CHYSEL sequence (pHR5640), or (d) an HIV construct with mCherry only as a negative control (pHR5461). In transfections (b) and (c), CMV-Rev (pHR30) was included to permit expression from the gag transcript. Forty-eight hours after transfection, cells were harvested and lysed. A Western blot was performed with extracts of the cells using primary antibodies to eGFP (Santa Cruz) and p24 (purified from Hybridoma 183-H12-5C NIH AIDS Reagent Program, Division of AIDS, NIAID, NIH, catalog number 1513[1]) as well as fluorescent secondary antibodies (LI-COR Biosciences). Imaging was performed using an Odyssey infrared imager (LI-COR Biosciences).

*Transfection-based functional assays.*

Except as noted, in transfection assays $8x10^5$ 293T/17 cells were transfected with 1000 ng of the HIV-derived construct and 100 ng of a Rev-containing construct (either pMSCV-Rev-IRES-fluorescent protein or CMV-Rev) in each well of a 12-well plate. Simultaneously, 293T/17 cells in different wells were transfected with constructs expressing individual fluorescent proteins to permit flow cytometry color compensation. At least one well of cells plated on the same day and not transfected with any construct was reserved as a control for flow cytometry gating purposes. 293T/17 cells were maintained prior to transfection in Iscove's modified Dulbecco's medium (IMDM) supplemented with 10% bovine calf serum (BCS) and gentamicin. One hour prior to transfection, cell medium was changed to IMDM+5% BCS.

Cells were incubated at 37°C for 24 hours after transfection. After incubation, 293T/17 cells were trypsinized and then suspended in phosphate buffered saline with 10% BCS to prevent further digestion. Suspended cells were transferred to a new tube, pelleted by centrifugation at 380 RCF for five minutes to remove the PBS+BCS, and finally resuspended in PBS alone.

Flow cytometry was performed using the Attune NxT flow cytometer with autosampler attachment (Thermo Fischer Scientific). Data acquisition was performed on the Attune NxT software package using the following channels when applicable:

| Laser lines | 488 | 405 | 561 |
|---|---|---|---|
| Emission filters | 530/30 | 440/50 | 620/15 |
| Fluorescent protein | eGFP | TagBFP | mCherry |

*Small molecule Rev-RRE pathway inhibitor assay.*

To apply inhibitor compound 103833 to cells, 10 uL of 103833-containing DMSO solution was dissolved in 1 mL cell culture medium (IMDM, 10% BCS, gentamicin) such that the final concentration of 103833 in the culture medium was 15 uM, 5 uM, 2.5 uM, 1 uM, 0.5 uM, or 0 uM. A total of 4 x $10^5$ 293T/17 cells were plated in each well of a 12 well dish in each concentration of 103833 cell culture medium. Twenty-four hours after plating, the 103833-containing medium was removed and replaced with IMDM+5% BCS. Each well of cells was transfected with one of two combinations of plasmids using the PEI method: 1000 ng pNL4-3(eGFP)(NL4-3 RRE)(mCherry) plus 100 ng pMSCV-NL4-3 Rev-IRES-TagBFP or 1000 ng pNL4-3(eGFP)(MPMV CTE)(mCherry) plus 133 ng CMV-Nxf1 plus 33 ng CMV-NxT1. Five hours after transfection, the 103833-free medium was removed from each culture and replaced with the original concentration of 103833 containing medium. Flow cytometry was performed 24 hours after transfection.

*Packaging and titering HIV and MSCV constructs.*

To package the HIV-derived construct pNL4-3(eGFP)(NL4-3 RRE)(TagBFP), a total of 3.5x$10^6$ 293T/17 cells were plated in a 10 cm dish. Twenty-four hours after plating, the cell culture was transfected with 15 ug pNL4-3(eGFP)(NL4-3 RRE)(TagBFP), 12.84 ug psPAX2, and 2.54 ug pMD2.G. Ninety-six hours after transfection, culture supernatant was removed, centrifuged at 380 RCF for five minutes to remove suspended debris, and filtered using a PVDF 0.45 uM syringe filter. Cell free medium was aliquoted and frozen at -80° C for future use.

To package the MSCV-derived constructs of the form pMSCV-Rev-IRES-mCherry carrying the NL4-3, 8-G, or 9-G Revs, a total of 3.5x$10^6$ 293T/17 cells were plated in a 10 cm dish. Twenty-four hours after plating, the cell culture was transfected with 15 ug pMSCV-Rev-IRES-mCherry, 12.84 ug pHIT60-CMV-GagPol[2], and 2.54 ug pMD2.G. Ninety-six hours after transfection, culture supernatant was removed, processed, aliquoted, and frozen at -80° C as above.

To determine the vector titer in the resulting stock solutions, CEM-SS cells were transduced with variable volumes of stock and the rate of successful transduction was determined by flow cytometry. Transductions were performed in duplicate. CEM-SS cells were suspended in PBS containing polybrene (final concentration of 10 ug/mL) and plated in 24 well plates at a concentration of $1x10^6$ cells per well. Stock vector-containing medium was thawed at 37°C and serial dilutions were made using IMDM+10% BCS+gentamicin as the diluent. Dilutions of virus stock were then added to the cells. The final volume of medium in each well was 1 mL, and wells contained either 200 uL, 20 uL, or 2 uL of vector stock solution. Cell cultures were returned to the incubator immediately after addition of viral stocks. Four hours after the addition of viral stocks, cells were pelleted by centrifugation and then replated on 6 well plates in 3 mL RPMI+10%FBS+gentamicin. Seventy-two hours after transduction, cells were collected by centrifugation, resuspended in PBS, and flow cytometry was performed. Cells that fluoresced with eGFP or TagBFP were scored as positive for pNL4-3(eGFP)(NL4-3 RRE)(TagBFP) and cells that fluoresced with mCherry were scored as positive for one of the pMSCV-Rev-IRES-mCherry constructs. Vector titer in the stock solution was calculated using the equation:

$$\frac{(Number\ of\ cells\ transduced \times Percent\ fluorescent)}{Volume\ of\ stock\ solution\ (mL)}$$

*Assay of Rev activity in transduced cells.*

To perform the transduction-level activity assay, CEM-SS cells were collected by centrifugation and split to plate $1 \times 10^5$ cells in each well of a 24 well plate in PBS containing polybrene at a final concentration of 10 ug/mL. Stocks of pNL4-3(eGFP)(NLR-3 RRE)(TagBFP), pMSCV-NL4-3 Rev-IRES-mCherry, pMSCV-8-G Rev-IRES-mCherry, and pMSCV-9-G Rev-IRES-mCherry were thawed and diluted in IMDM with 10% BCS to achieve a predicted multiplicity of infection of 0.1 for each construct. The diluted vectors were added to the CEM-SS cultures either individually or in combinations of the RRE-containing construct with each Rev-containing construct.

After the addition of the viral vectors to the cells, each plate was subjected to spinoculation by centrifugation at 380 RCF for two hours at 25°C[3, 4]. The plates were then placed in an incubator for two hours at 37°C. Next, the suspended cells and medium were collected and centrifuged at 180 RCF for seven minutes to pellet the cells. The medium was removed from each cell pellet, and cells were resuspended in 200 uL RPMI with 10%FBS and gentamicin and then transferred to a 96-well U-bottom plate. The resuspended cells were returned to the 37°C incubator. Seventy-two hours after transduction, cells were harvested, pelleted by centrifugation, washed, and resuspended in PBS. Then, flow cytometry was performed on the cell suspensions as above. Functional activity was determined by gating a population of cells successfully transduced with both constructs and calculating the ratio of eGFP:TagBFP MFI (see Figure 2 panel e).

*Creation of T-cell line containing the integrated two color reporter construct.*

An HIV-derived construct was created of the form pNL4-3(eGFP)(NL4-3 RRE)(mCherry) with an additional deletion in *vpr* to prevent expression of that viral protein (pHR5604). The construct was packaged and VSV-G pseudotyped as above, then CEM-SS cells

CEM-SS cells were transduced using DEAE-dextran. The bulk culture was split once, then a limiting dilution was performed and single cells were plated on a 96 well plate to generate monoclonal populations. The dilution plates were incubated for two weeks, then clones expressing the HIV-derived construct were identified by looking for mCherry expression via florescent microscopy.

Inverse PCR was performed on selected clones to ensure the presence of a single integrated copy of the HIV-derived construct and to locate the site of integration. Genomic DNA was prepared from each clone, cut with SpeI, and then self-ligated to form circular fragments. Nested PCR was performed on the bulk ligation reaction using forward primers located within *gag* and reverse primers within the 5' LTR. Sanger sequencing was performed on the resulting PCR product using the inner primers and the sequence was analyzed using Geneious R8 (Biomatters, Auckland, New Zealand) to identify HIV and cellular sequences. The cellular sequences were mapped to the GRCh38/hg38 assembly using the UCSC Genome Browser (http://genome.ucsc.edu/)[5]. Primers used in this procedure:

| **Directionality** | **Nested** | **Oligo number** | **Sequence (5'-3')** |
|---|---|---|---|
| Forward | Outer | 3757 | GGTCAGCCAAAATTACCCTATAGTG |
| Forward | Inner | 3759 | TGTTAAAAGAGACCATCAATGAGGAAG |
| Reverse | Outer | 3765 | CACGTGATGAAATGCTAGGCG |
| Reverse | Inner | 3766 | TCAGTGGATATCTGACCCCTGG |

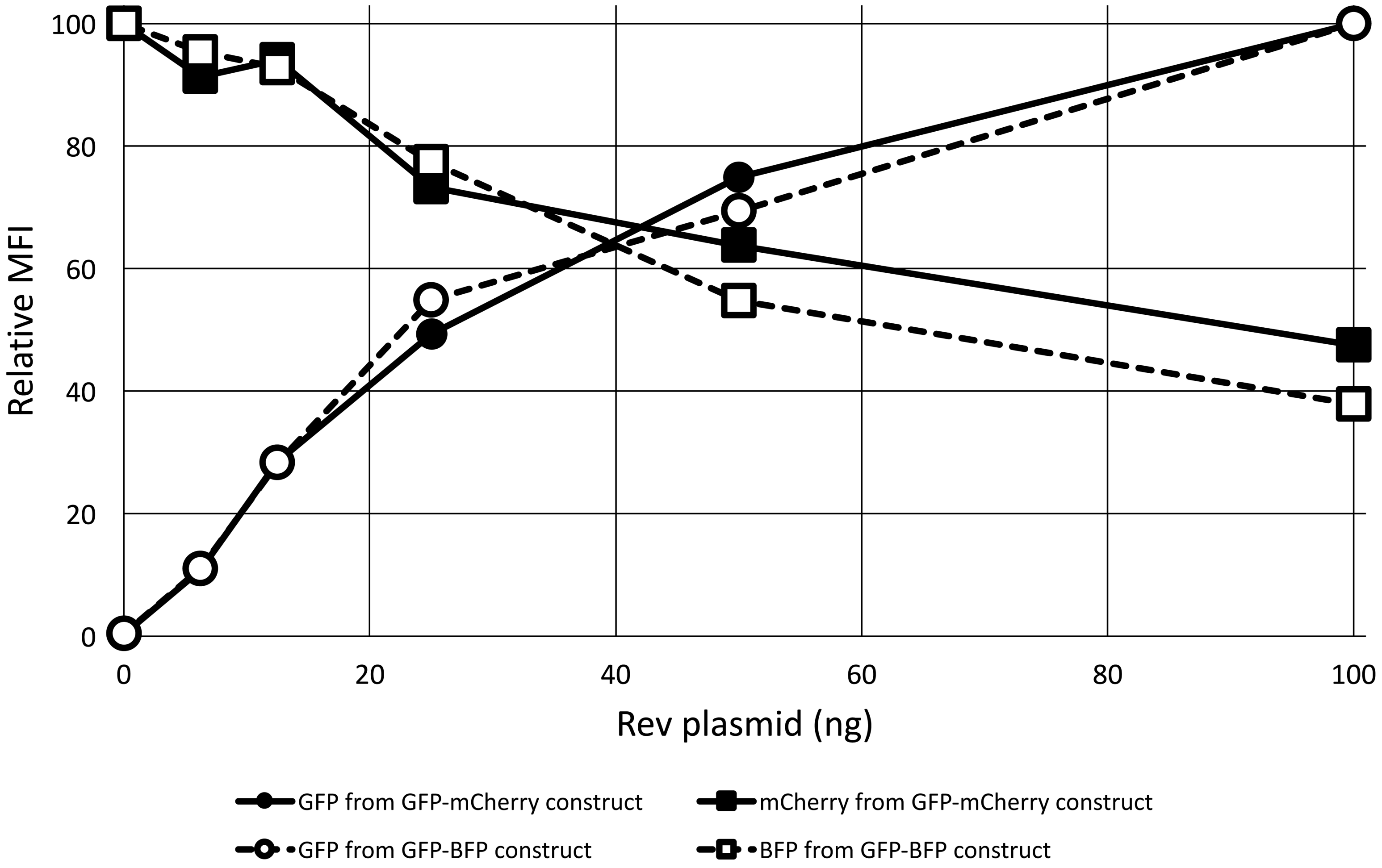


Figure S1. Independent measurement of fluorescent markers from Rev-dependent and Rev-independent transcripts. 293T/17 cells were transfected with either eGFP-mCherry or eGFP-TagBFP HIV-derived constructs along with varying amounts of CMV-Rev plasmid, as in Figure 3. For each HIV-derived construct and each experimental condition, the mean fluorescence intensity of both the Rev-independent (TagBFP or mCherry) and Rev-dependent (eGFP) fluorescent markers was measured. The maximum MFI for each marker on each transcript was set as 100, and the other values for that marker were normalized accordingly.

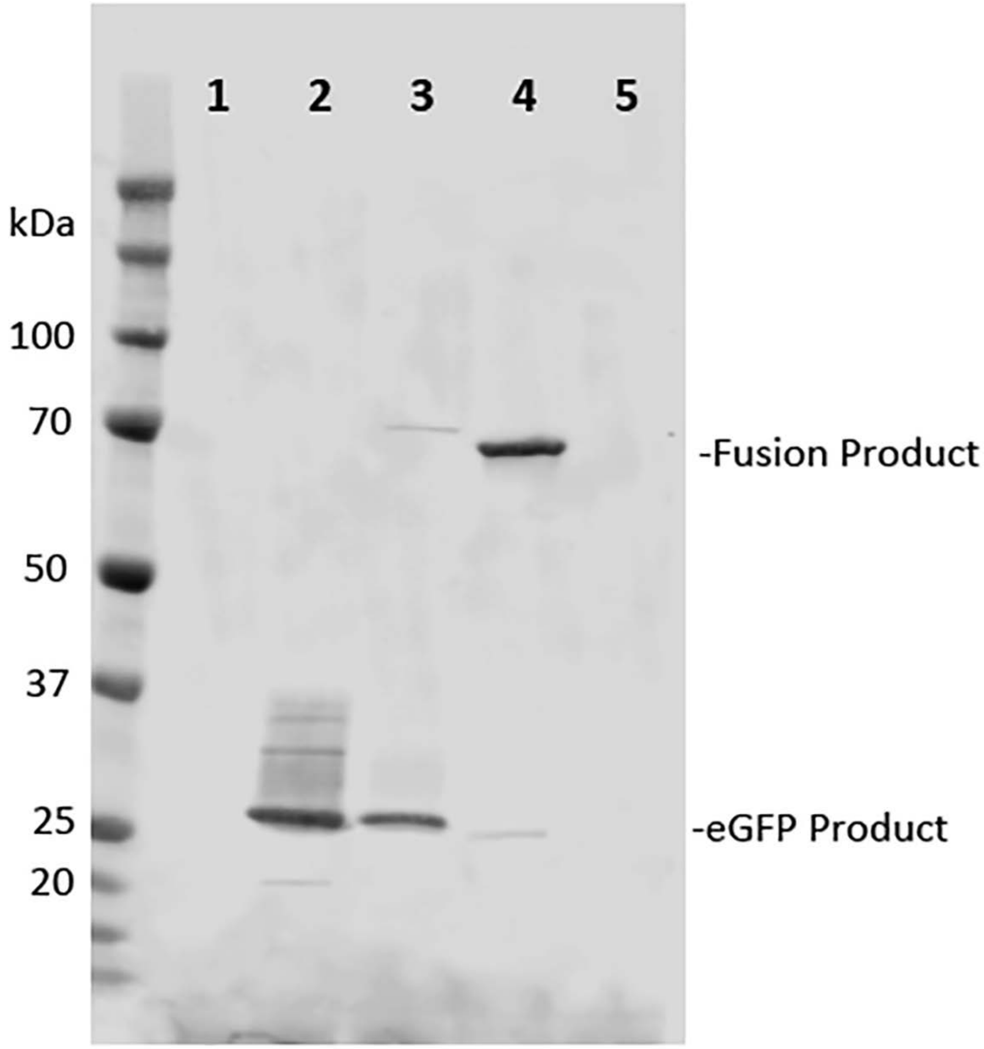


Figure S2. The P2A CHYSEL sequence allows the production of separate Gag and eGFP proteins. To produce a fluorescent signal in a Rev-dependent fashion, *gag* was truncated and a cassette was inserted in frame consisting of a P2A CHYSEL sequence and the eGFP gene. The CHYSEL was expected to result in the production of two separate polypeptides – a Gag fragment and a free eGFP. Western blotting on of an SDS-polyacrylamide gel was performed to demonstrate that this cleavage was efficient. The entire uncropped blot is shown. To produce this blot, 293T cells were either directly lysed (lane 1) and analyzed, or lysed and analyzed after transfection with constructs producing eGFP alone (lane 2), an HIV vector similar to Figure 1A with Gag-CHYSEL-eGFP (lane 3), an HIV vector the with a Gag-eGFP fusion sequence without a CHYSEL (lane 4), or an HIV vector producing only mCherry (lane 5). Blotting was performed using an antibody directed against eGFP. Free eGFP is seen in lanes 2 and 3. The 70 kDa band in lane 4 corresponds to the Gag-eGFP fusion product.
